# Recommendations for Bioinformatics in Clinical Practice

**DOI:** 10.1101/2024.11.23.624993

**Authors:** Ksenia Lavrichenko, Emilie Sofie Engdal, Rasmus L. Marvig, Anders Jemt, Jone Marius Vignes, Henrikki Almusa, Kristine Bilgrav Saether, Eiríkur Briem, Eva Caceres, Edda María Elvarsdóttir, Magnús Halldór Gíslason, Maria K. Haanpää, Viktor Henmyr, Ronja Hotakainen, Eevi Kaasinen, Roan Kanninga, Sofia Khan, Mary Gertrude Lie-Nielsen, Majbritt Busk Madsen, Niklas Mähler, Khurram Maqbool, Ramprasad Neethiraj, Karl Nyrén, Minna Paavola, Peter Pruisscher, Ying Sheng, Ashish Kumar Singh, Aashish Srivastava, Thomas K. Stautland, Daniel T. Andreasen, Esmee ten Berk de Boer, Søren Vang, Valtteri Wirta, Frederik Otzen Bagger

**Affiliations:** Oslo University Hospital; Rigshospitalet; Karolinska University Hospital; Karolinska Institutet; Haukeland University Hospital; Institute for Molecular Medicine Finland; National University Hospital of Iceland; Turku University Hospital; Skåne University Hospital; Helsinki University Hospital; University of Helsinki; University Medical Center Groningen; University Hospital of Umeå; KTH Royal Institute of Technology; St Olav’s University Hospital; Aarhus University Hospital

## Abstract

Next Generation Sequencing (NGS) is increasingly used in clinical diagnostics, largely driven by the success and robustness of Whole Genome Sequencing (WGS). Whereas updated guidelines exist for how to interpret and report on variants that are identified from NGS using bioinformatics pipelines, there is a need for standardised bioinformatics practices for diagnostics to ensure clinical consensus, accuracy, reproducibility and comparability of the results. This article presents consensus recommendations developed by 13 clinical bioinformatics units taking part in the Nordic Alliance for Clinical Genomics (NACG), by expert bioinformaticians working in clinical production. The recommendations are based on clinical practice and focus on analysis types, test and validation, standardisation and accreditation, as well as core competencies and technical management required for clinical bioinformatics operations.

Key recommendations include adopting the hg38 genome build as the reference and a standard set of recommended analyses, including the use of multiple tools for structural variant (SV) calling and in-house data sets for filtering recurrent calls. Clinical bioinformatics production should operate under the ISO 15189 standard, utilising off-grid clinical-grade high-performance computing systems, standardised file formats, and strict code version control. Containerized software containers or environment management systems are needed to ensure reproducibility.

Pipelines should be rigorously documented and tested for accuracy and reproducibility, minimally covering unit, integration, and end-to-end testing. Standard truth sets such as GIAB and SEQC2 for germline and somatic variant calling, respectively, should be supplemented by recall testing of previously validated clinical cases. Data integrity must be verified using file hashing, and sample identity should be checked via sample fingerprinting and genetically inferred identification markers such as sex and relatedness.

Finally, clinical bioinformatics teams should encompass diverse skills, including software development, data management, quality assurance, and domain expertise in human genetics. These recommendations provide a consensus framework for standardising bioinformatics practices across clinical WGS applications and can serve as a practical guide to facilities that are new to large-scale sequencing-based diagnostics, or as a reference for those who already run high-volume clinical production using NGS.

## Introduction

In the field of clinical variant interpretation and classification, there are best practice recommendations available, such as ACMG guidelines (Richards et al., 2015), and ACGS guidelines (ACGS, 2020). However, these guidelines rely heavily on the bioinformatics pipelines where the variants are called. In 2018, Roy et al. presented consensus recommendations for the validation of clinical NGS bioinformatics pipelines (Roy et al., 2018). Since then, the requirements in the field have grown, notably in the throughput of samples, and the size of the data. Both parameters are largely driven by the success of Whole Genome Sequencing (WGS) which has proven a major advantage over targeted gene panels or exome sequencing, both of germline cells for diagnostics of hereditary diseases (Splinter et al., 2018), (Genomes Project Pilot Investigators et al., 2021), (Stranneheim et al., 2021) and for identification of treatment targets in somatic mutations for cancer diagnostics (Sosinsky et al., 2024), (Hodder et al., 2024). Therefore, sequencing-based diagnostics units now handle a growing number of samples, and for each of these, more data is generated. Moreover, there has been an increase in expectations regarding turnaround time (TAT), quality standards, and performance evaluation within an operating clinical bioinformatics unit. An increase in knowledge on the optimal analysis pipeline, as well as new diagnostic scores and markers also means a higher number of expected analysis outputs from the same sample, and it also calls for individually tailored analysis targeted towards specific patient groups (Bagger et al., 2024). Therefore, a classic bioinformatics core facility providing support for medical research (Chicco & Jurman, 2023) is now a very different entity compared to units supporting production-scale sequencing analysis for diagnostics, even if there can be some competencies that overlap. The development towards professional organisation of large-scale automated continuous production in the field of clinical bioinformatics has only been accelerated with increasingly larger capacity sequencers that require a high sample influx for cost-effective production and a low per-sample price (Eisenstein, 2023), and acceptable TAT for urgent samples.

Here, we report on current practice in 13 clinical bioinformatics units in the healthcare systems of the Nordic (and Northern Europe) countries and provide a set of general recommendations for clinical bioinformatics at scale. We will cover the processes from raw data output from the sequencer to data processing and analysis steps until interpretable information, usually a file with variants, or a clinical score.

The recommendations are based on a survey, a workshop, and final approval by members of Nordic Alliance for Clinical Genomics present at the Clinical Bioinformatics workshop in Helsinki 2023 (NACG 14th, 2023). By design, only recommendations that could achieve unanimous votes have passed, and as such all authors have the right to veto any recommendation. In brief, a summary of the survey provided the starting point for a presentation at a plenary session where first the most radical interpretation of a recommendation was presented, and subsequently, soften the language or moderate the statements until a full agreement could be reached, or the recommendation was rejected. Here we give recommendations that are based on clinical practice of at least two sites. In practice, we found that most sites follow most of the recommendations.

The scope of our survey and workshop was primarily Next-Generation Sequencing (NGS), acknowledging that there may be differences between tools and pipelines for targeted panels, exome, and whole-genome sequencing (WGS). We had as an aim to present a perspective that can be generalised to all these settings, and when differences were incompatible we would prioritise emphasis on WGS.

We acknowledge the relevance of sequencing platforms like long-/short-read and different vendors and technologies, we decided a priori not to specifically address platform-specific recommendations, but merely record the platforms in use.

All authors and participants in the survey are running clinical diagnostics production and our recommendations aim to serve as a practical guide for hospitals or facilities new to large-scale sequencing-based diagnostics, or as a reference for inspiration or discussion for those who already run high-volume clinical production using NGS.

## Methods

Recommendations were based on a survey (Supplementary S1) sent to NACG members participating in the workshop for Bioinformatics in Clinical Practice held in Helsinki 28-29 September 2023. Participants were, in the invitation, specifically asked to compile answers from their unit, and as such represent opinions from the team. All statistics are based on the survey, and members have been asked to confirm that their views are represented whenever free-text information had to be summarised. Based on the answers a series of statements was discussed at the workshop, where the most extreme statement that could be devised from the questionnaire was put forth first, and if concerns were raised by anyone, a less extreme alternative was put forth, until consensus could be found. Every member had veto right on any statement. Three topics were planned for more detailed active discussions; participants were divided into three teams that were rotated three times to visit all topics, with a static session chair. The session chair summarised the discussions into recommendations that were put to vote in a final plenary session.

Participants were asked to support recommendations that they, based on their expertise, would recommend for clinical bioinformatics production. Therefore not all sites have a production that follows all recommendations, but by design, no recommendation is given that is not in production in at least two sites. All sites have been promised anonymity in their answers in this publication, to ensure that we can maintain the honest and open discussion that is at the heart of NACG.

## Results and discussion

Based on a survey and a workshop in Helsinki in September 2023, with members of NACG we arrived at 16 recommendations for clinical bioinformatics production (Table 1), as well as summary statistics on sample numbers, methods and implemented standards at each site (Fig.1 and Supplementary Table T1)

**Table 1:** Recommendations for sequencing-based clinical bioinformatics. Abbreviations are SNV: Single Nucleotide Variant; CNV: Copy Number Variant; SV: Structural Variants; STR: Short Tandem Repeat Expansions; LOH: Loss of Heterozygosity; PRS: Polygenic Risk Score; TMB: Tumour Mutational Burden; HRD: Homologous Recombination Deficiency; MSI: Microsatellite Instability; HPC: High Performance Computing, GIAB: Genome in a Bottle; SOP: Standard Operating Procedure.

|  |
| --- |
| 1. Genome build Hg38 is the reference for alignment. |
| 2. A recommended standard set of analyses: SNV, CNV, SV, STR, LOH, variant annotation, PRS (optional), and for cancer, TMB, HRD, MSI. |
| 3. Several tools are needed in combination for calling structural variants. |
| 4. Structural variants must be filtered using a tool-specific matched in-house dataset, to filter common variants and false positives variant calls. |
| 5. Use reliable air-gapped clinical production-grade HPC and IT systems. |
| 6. Clinical bioinformatics production should operate under ISO15189. |
| 7. Standardised file formats and terminologies should be used. |
| 8. Pipelines should be well documented, and tested for both accuracy and reproducibility against predefined acceptance criteria. |
| 9. Production code must be subjected to manual review and testing. |
| 10. Computer code and documentation must be developed and managed under strict version control in a git-tracked system. |
| 11. Validation using standard truth sets, GIAB (germline) and SEQC2 (somatic), should be accompanied by a recall-test of previous clinical cases, preferably from orthogonal methods. |
| 12. Pipelines should be tested at the levels of unit, integration, system, IT performance, and end-to-end tests. |
| 13. Integrity of data must be verified using file hashing (e.g. MD5 or sha1). |
| 14. Identity of sample should be verified via inference of identifying traits and checks for relatedness between samples (sample fingerprinting tests). |
| 15. Software should be encapsulated in containers or conda environments. |
| 16. Clinical bioinformatics teams must be able to attract a combined skill set covering software development, data management, quality assurance, and domain knowledge in human genetics. |

### NACG community

**Figure 1.**
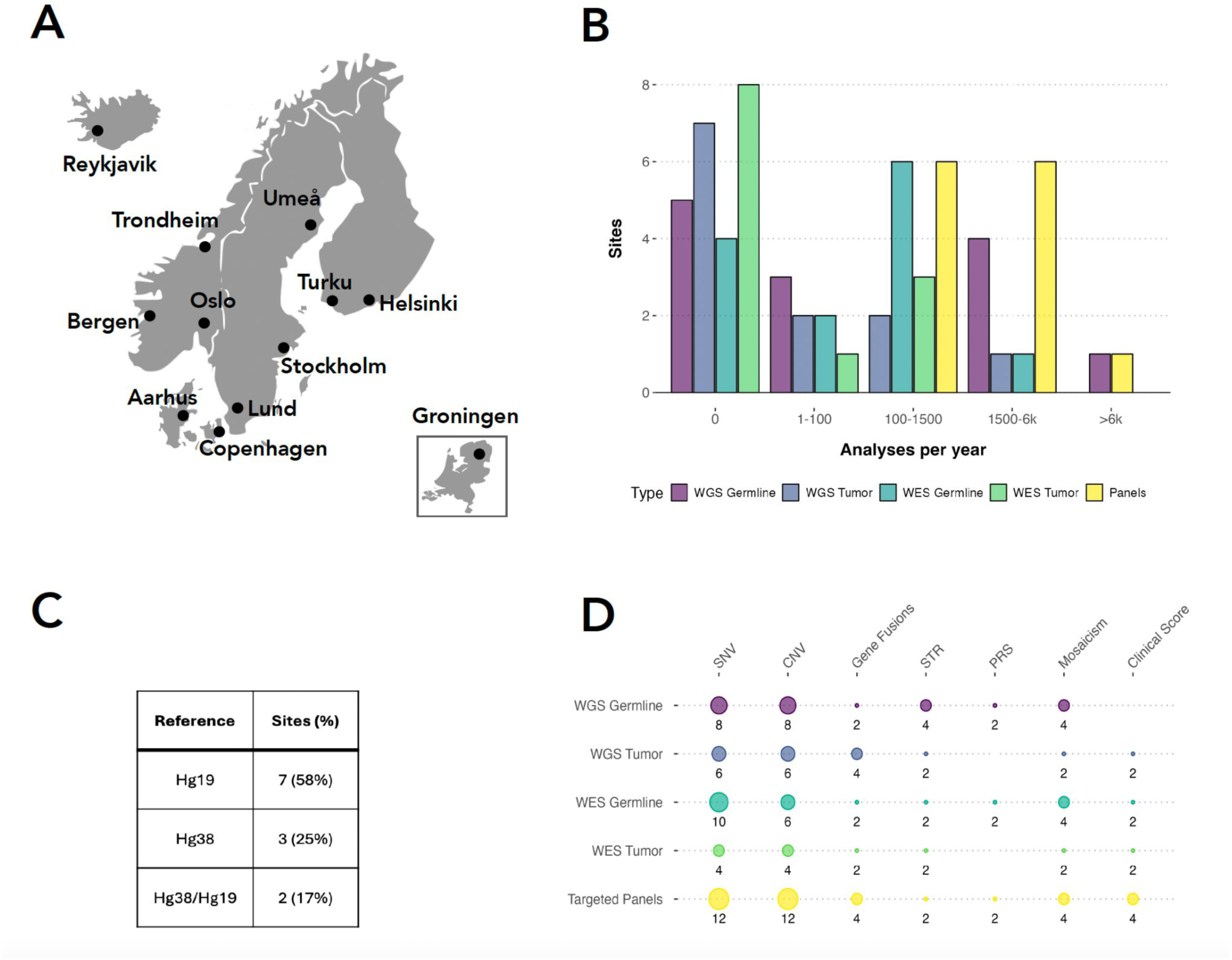
Overview of the NACG community. A. Geographic distribution of NACG sites: a map illustrating the 12 member sites of NACG network and one guest site from The Netherlands. B. Annual analysis volume per site: a bar chart depicting the volume of analyses performed per year across NACG sites. The x-axis shows four categories of analysis volume, while the y-axis indicates the number of sites (out of 13) in each category. Different colours represent various types of analyses, as indicated in the legend. C. Genome reference build usage: a summary of genome reference build statistics across the 13 participant sites. D. Details of pipelines in clinical production on a matrix plot: the y-axis shows the type of analysis, and the x-axis shows the type of variant or metric reported. Bubble size reflects the number of sites in each intersection category. The colour legend is the same as in panel B.

### A recommended set of analyses for clinical NGS

The set of recommended analysis for clinical production of WGS is in line with previous reports (Bagger et al., 2024), (Stranneheim et al., 2021), (Kobren et al., 2021), with some optional points specific on whether the facility is performing cancer analysis or not. Automated quality assurance is usually handled partially, or fully, within the analysis pipeline; it is a multifaceted task influencing all aspects of production, which will be handled in later sections.

Whereas there is a core set of operations that all facilities have to perform, there are - even when using the same software or tools - a multitude of parameters that should be set and optimised differently depending on the purpose and nature of sequencing data. In the following, we will stay at the level of describing a core set of analyses that we recommend for NGS based diagnostics, see below:

- **De-multiplexing** of raw sequencing output to disentangle pooled samples (BCL to FASTQ),
- **Alignment** of sequencing reads to a reference genome (FASTQ to BAM):
- **Variant calling** (BAM to VCF)

- SNVs and small insertions and deletions (indels)
- CNVs (deletions and duplications)
- SVs including insertions, inversions, translocations and complex rearrangements
- STRs
- LOH: loss of heterozygosity regions (indication of UPD, uniparental disomy)
- Mitochondrial SNVs and indels
- **Variant annotation** (VCF to annotated VCF).

Of note, SNVs and indels (<50 base pairs) are often handled and conceptualised together because the same tools can output both. Inversely, copy number variants (CNV) and short tandem repeat expansions (STR) are often handled separately from other structural variations (SV) because tools and methods, at least historically, were dedicated to either one type, or the other. Mitochondrial variant calling also benefits from a tailored approach.

Variant annotation was not part of the scope for these recommendations. It is a large and complex task and requires significant maintenance to keep the annotation data sources relevant and updated. By some units the choice of annotation is considered the domain of the clinical interpretation, and by others it is handled in collaboration with the bioinformatics unit - also depending on the graphical user interface software used for variant interpretation and classification. There are also differences as to whether to use soft-filter (just add information in the form of tags to the VCF-file), or to hard-filter (remove variants from the VCF-file) that do not pass a given criteria. Similarly, some units approach the variant annotation task as a variant prioritisation problem (ranking of variants from likely clinically relevant to likely irrelevant), and others by providing a fixed list for interpretation (often in the form of predefined gene panels).

#### Optional analyses

- Microsatellite instability (MSI): MSI analysis assesses mutations in microsatellite regions to identify DNA mismatch repair defects, used for guiding immunotherapy treatment in cancers.
- Homologous recombination deficiency (HRD): HRD testing evaluates the integrity of homologous recombination repair pathways predicting response to PARP inhibitors in cancers, particularly ovarian and breast cancer.
- Tumour mutational burden (TMB): TMB quantifies the number of somatic tumour mutations. This is used as a proxy for the production of neoantigens that could trigger an immune response, and TMB thus identifies patients likely to benefit from immunotherapy.
- Polygenic risk scores (PRS): PRS estimates an individual’s genetic predisposition to complex diseases by aggregating effects of multiple genetic variants. Only one site reports to have this in production and it may be pending further validation and standardisation.

### Genome build hg38 as a reference

Not all sites have yet switched from GRCh37/hg19 to GRCh38/hg38 (see Fig. 1C), but this transition is a recommendation, despite the significant transition cost. The hg38 build has reduced gaps, improved mappability and variant calling, and corrects several clinically relevant errors present in the hg19 reference (Li et al., 2021). In addition some of the important public reference data repositories continue their releases only in GRCh38/hg38 build - such as gnomAD v4 (gnomAD, 2023).

The transition, however, touches upon a large number of dependencies both in the bioinformatics processing, variant annotation and legacy data that potentially needs to be re-analyzed. Using automated tools to lift local accumulated variant classification and other manual annotation is not optimal, and usually means that any accumulated manual knowledge is not reliable for automated translation between the two different genome references (Pan et al., 2019). The public annotation databases, tools and pipelines, relevant for clinical production, now all support hg38, with the exception of the panel-based TSO500 Illumina DRAGEN pipeline (Illumina DRAGEN TSO500, 2023), that is in production at several sites.

### Several tools are needed in combination for calling of structural variants

There are several strategies for calling structural variants, including models based on read depth quantification, assessment of split read mapping, distant or mis-oriented mapping of read-pairs, and de novo assembly (Gabrielaite et al., 2021). All sites experience and agree that all tools for calling structural variants have their caveats, and that a combination of tools and strategies are needed, which is also what most sites have in production. There are different approaches to how to integrate the output from several callers which could be either a consensus of calls (intersect), a merge of calls (union) or a weighted output that includes uncertainty measures, either at the variant level, or the level of the tool (e.g. we always include calls from X, but only from Y, if there is agreement with Z).

#### False positive filter

To control for the large amount of false positives all sites use an in-house database of previously called variants, containing the frequency of how many times a certain structural variant has been called before. This database, sometimes referred to as a false-positive-filter (FPF) will consist of a mix of false positive calls and real but frequent variants, both of which are uninteresting for identification of clinically relevant rare variants (Eisfeldt et al., 2017). Given the high number of false positives from any SV calling tool (Gabrielaite et al., 2021) determined by the specific strategy for calling the variants, the background database must be reconstructed at every update of the pipeline that significantly affects the calls, be it from a new version of the tool, or change in upstream processing. The practical experience is also that changes in wet-lab library preparation kit, the sequencing chemistry, and sequencer also have a sizable impact on the nature of false calls, just like the extent of familial ancestry between the patient and the samples in the FPF will impact the effectiveness of the filter (Eisfeldt et al., 2020).

#### Comparing structural variants

In order to use a FPF it is necessary to be able to compare variants from a patient to a database. It is a non-trivial task to compare two or more structural variants because the precise break-ends or structural variant class is often not called with precision. Therefore tools like SVDB (https://github.com/J35P312, 2019) can be used to set criterias for overlap both in terms of distance between the break-ends and the required percentage overlap.

### Accreditation and quality standards

It is of importance for clinical laboratories to ensure trust in the quality of the performed work. A way to achieve this is to obtain accreditation by authoritative bodies (national accreditation bodies) to certify that the laboratory operates according to a certain scheme, e.g. a standard defined by International Standards Organization (ISO). Specifically, the ISO 15189 standard (newest version is ISO 15189:2022) on “Medical laboratories - Requirements for quality and competence” is relevant for clinical laboratories or units that perform medical tests based on patient material, e.g., extensive genetic analyses. All participants performed analyses according to the requirements of either ISO 15189 or ISO 17025 standards (12 and 1, respectively). The difference between them is that ISO 17025 is a general standard for testing and calibration laboratories, while ISO 15189 is specific to medical laboratories. ISO 17025 serves as a normative reference for ISO 15189.

The accreditation is obtained for specific analyses, and it may not be feasible for a laboratory to have all analyses accredited, e.g., due to cost associated with accreditation, low volume of samples for the given analysis, frequent changes in procedures, or large heterogeneity in the procedures required for different sample types.

The scope of an accredited analysis is defined by the accredited laboratory, and as such accredited analyses are, while they may be similar, often not equivalent across accredited laboratories. E.g., two separate analyses in one laboratory may correspond to a single analysis in another laboratory. Accordingly, the mere count of analyses each laboratory has accredited does not translate into to which extent the accredited analyses cover the possible spectrum of genetic tests. Instead, we asked the participants for the aims of the bioinformatics pipelines that were used in accredited analyses. The accredited analyses across the 13 participants included bioinformatics pipelines aimed for one or more of the following analyses: call of germline small variants, copy number variants, and structural variants in whole genome sequencing data from DNA; somatic small variants in whole genome sequencing data from DNA; gene fusion variants in sequencing data from total RNA; and small germline variants and copy number variants in sequencing data from DNA gene panels (Supplementary figure/table S1).

Quality schemes, e.g., like the ISO 15189 standard, set requirements to cover a wide range of aspects that influence the quality of the work. While we do not make recommendations for all such aspects here, the participants agreed that methods within clinical bioinformatics are often not standardised, highly customised per laboratory, and used for a scope that is not covered by the validation performed by the provider. Also, there is a lack of relevant external quality assessment programmes. Accordingly, all laboratories recommended special attention to, that bioinformatics are rarely externally validated for scopes relevant to clinical usage, but requires validation by the user.

All participants recommended to work according to, and ideally to be accredited according to, an ISO 15189-equivalent quality management system. Furthermore, a future implication is that work performed according to ISO 15189 standards fulfils European Union regulations 2017/745 (Medical Devices Regulation) and 2017/746 (In Vitro Diagnostic Medical Devices Regulation (Regulation (EU) 2017/746, 2017); requirements for laboratory-developed devices (e.g. clinical bioinformatics pipelines) for which an equivalent CE-marked device is not available ((MDCG), 2023).

### Safety, quality and compliance

In clinical bioinformatics a robust quality management system is a requirement, fully in line with any other diagnostics procedure, and very different from research- or project-driven bioinformatics. Importantly, clinical diagnostic production relies on the ability to process the samples continuously, rather than by manual handling of batches. This entails the need for a high level of automation and systems integration, and both data and operations must be designed and conceptualised as flows, rather than units handled successively. This additionally removes many sources of human errors, and allows for scalable production. Automated quality procedures must balance the need for every single patient to receive an answer (it is not a viable solution to simply discard a large fraction of the samples to get a high quality dataset) and the detrimental effects that a wrong answer could have for a patient. The latter is further accentuated by the fact that germline genomic data is a lasting resource for later diagnostics and general reuse by the patient, immediate family and society at large.

#### Personal identification

Patient information must be stored safely and be correct. Genomic information is as per definition both personal and sensitive in the European Union (GDPR recital 34), and additionally, healthcare information - by bioinformaticians sometimes referred to as clinical metadata - is needed to run bioinformatics pipelines and to analyse the results. In some instances, it becomes a balancing act between data security, which often presents itself as a form of inaccessibility and data barriers, and patient safety that will suffer, if the healthcare professionals do not have easy and operational access to the right information at the right time, or are forced to adopt either manual or unintended procedures to circumvent rigidly designed safety structures. In addition to high requirements for data handling and data flow, verification and validation must be in place to confirm data integrity in dataflows; all sites use and recommend automated check of hashes, e.g. MD5 og sha1, for data verification. Additionally, sample identity checks is a recommendation to detect swaps or mixing, and can be in the form of dual sequencing of samples in two different laboratory and data processing flows - one of which only needs to contain a small number of polymorphic sites - and subsequent relatedness check (Pedersen et al., 2020), or relatedness checks can be performed within samples from the same sequencing flowcell, or within a time window, and obviously between family members when a full trio is sequenced for diagnostics of inherited diseases. Other sample identity checks include assessment and control of data from sex chromosomes X and Y (Liu et al., 2022) and ancestry inference.

#### Quality control

There was no consensus on which parameters to use for automated quality control (QC). All units aim for either a mean or median sequencing coverage depth of >30-fold for germline WGS and 60-90-fold for somatic WGS, but little consensus exists regarding minimal coverage depth for panels or WES. Other scores used for quality assessment are Phred scores, estimated sequencing error rate (Q20 or Q30), allele depths ratio, Q pred scores, GC ratio, AT dropout and skewed allele depth ratio (<0.05). Thresholds for most of these quality scores are used in combination with a definition of a fraction of the genome that should pass the threshold, in order to account for hard-to-sequence regions of the genome, while still applying a stringent threshold. Several units report using soft thresholds or have intervals that do not necessarily stop the sample from being sent to the interpreting personnel, but flagging it with a warning. The tools reported to calculate and collect automated QC in production are: FastQC, MultiQC Picard, GATK, Illumina SAV, Qualimap, OmnomicsQ, MultiQC, Mosdepth, SamBamBa, and in-house tools. All unit reports that manual visual inspection of variants are performed during the variant interpretation by a person skilled in the art (e.g. geneticist, biologist or medical specialist) prior to drafting the clinical report.

**Figure 2.**
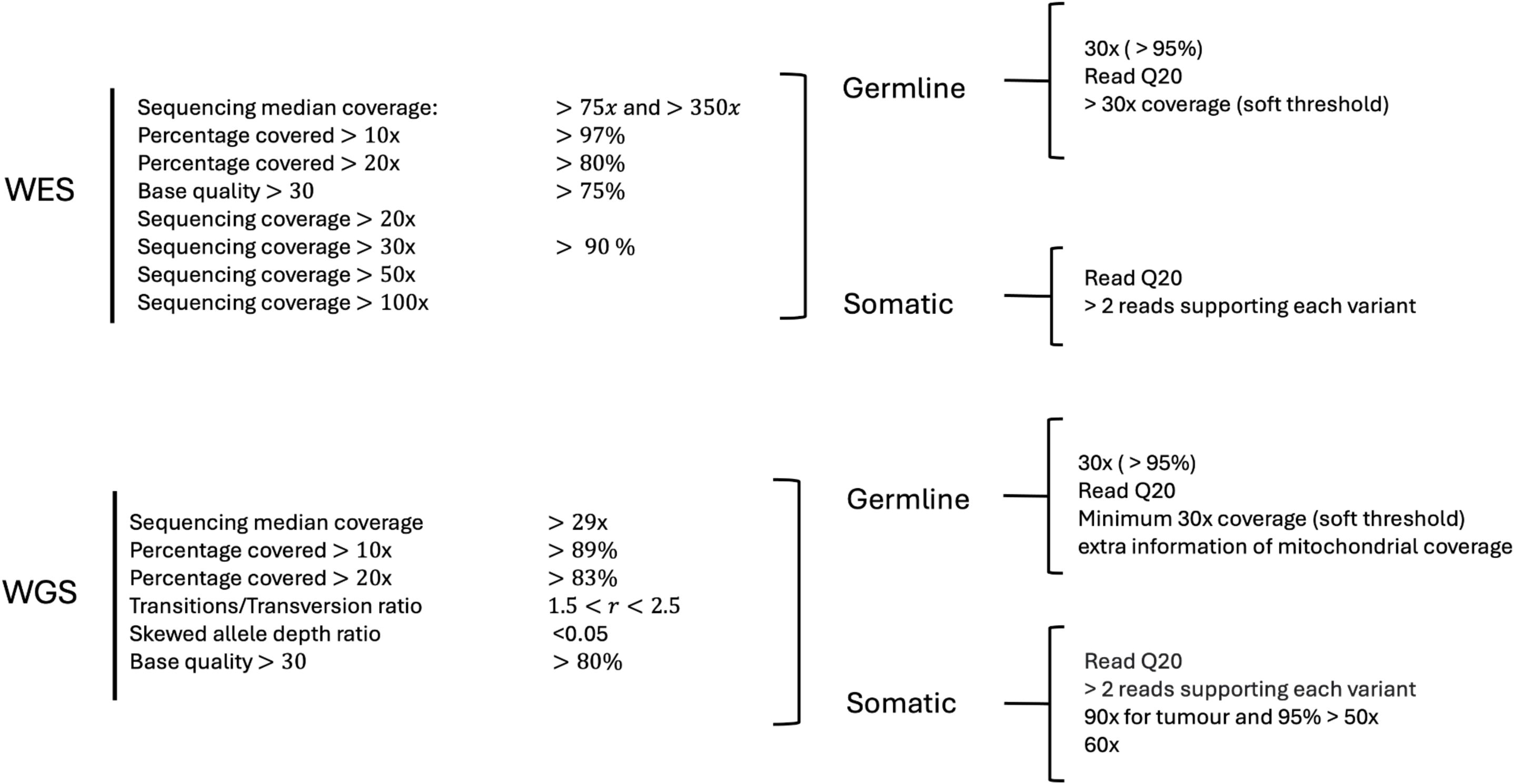
Summary of reported coverage and quality threshold used in clinical production. Consensus of minimal thresholds and quality metrics remains limited for targeted panels and WES, whereas a minimum sequence coverage depth is agreed upon across all units for both somatic and germline testing.

Coordination of quality management varies depending on the organisation or hospital that the unit operates in. In some cases, a dedicated team handles all quality-related tasks, also including accreditation and audits. Such dedicated teams may not include competences in bioinformatics or computational engineering, and since the available standards have been devised for medical laboratories, this can lead the bioinformatics team to adapt to practices more suited for wet lab. Alternatively, quality tasks may be managed by bioinformatics team members who also work in the field, but may have less experience in broad quality practices. Both approaches have their caveats, and the choice is often determined more by the available resources and competences at the site, rather than design. All sites agreed that accreditation and quality work is valuable, also because it helps in maintaining a quality and safety aware culture.

#### Version control

A fundamental part of accreditation according to the standard is documentation of procedures. To most bioinformatics or general software developers this is a natural part of good coding practices, and rarely needs fundamental changes in working routines. Use of git is recommended for collaborating on code as it makes it both easy and natural to enforce version control and support release management. The choice of platform is even between github (5/13, www.github.com) and gitlab (5/13, www.gitlab.com).

Broader description of the pipelines, the tools and the components, as well as test results, is necessary in addition to the code and user documentation. Versions of the software and pipelines, for example, must be clearly documented and accessible, not obscured within git tags, and available to anyone receiving output from the pipeline.

### Benchmarking, validation, and testing of bioinformatics tools and pipelines

#### Test data

An important aspect of ensuring high quality in production is the ability to quantify the results. Truth sets for test and validation comprise a set of sequencing data with the corresponding target - e.g. variants or diagnostics score - that is the desired result of an analysis. These should be designed to be representative of clinical samples and variants to be analyzed, and data should preferably have been sequenced in-house, to account for local technical biases. For common truth sets the biological material can be acquired for this purpose. There is recognition of the overreliance on the major public truth sets from the Genome in a Bottle Consortium (GIAB) (Jarvis et al., 2022), (Wagner et al., 2022), (Zook et al., 2020), (Krusche et al., 2019), (Zook et al., 2019), (Berner & Law, 2016) and The Sequencing Quality Control 2 (SEQC2) (Fang et al., 2021), which are often used both for training tools and for validation. This dual use introduces double dipping, and highlights the need for additional local truth sets to avoid overfitting and ensure robust performance, even if these additions are more difficult to work with. A dedicated test set to identify variants in hard-to-call clinically relevant genes (Wagner et al., 2022), can make the test more relevant and realistic, and adds variants not present in the standard GIAB truth set. There are also multiple External Quality Assessment (EQA) programmes (Hastings & Howell, 2010) with the same aim albeit often in a blinded validation that does not lend itself to further testing. All units use, in some form, previous samples to test and validate new pipeline releases, knowing that it carries a clear bias towards recall, at the expense of precision. Most treasured recall-sets are re-sequencing of samples where the target have been identified with orthogonal methods like SNP arrays, Sanger sequencing or Multiplex ligation-dependent probe amplification (MLPA) or even manual “eye-balling” of sequencing reads, because these cases alleviate the known and potentially monumental recall bias towards the approaches or algorithms used for generating the target in the truth.

Use of synthetically introduced variants using BAMsurgeon (Ewing et al., 2015) have previously been tested within NACG as a communal quality assessment effort (NACG 10th, 2021) but no unit uses this strategy routinely. One concern that arose from the test is that synthetically introduced variants fail to trigger thresholds that depend on the neighbourhood of the variant, e.g. identification of active regions, local realignment, or scores that comprises a genomic-spatial effect including read- and mapping-quality, as well unknown technical artefacts including editing the distance between sample and the reference genome.

For testing purposes several facilities use reduced datasets, e.g. data from only select chromosomes or otherwise truncated raw data, to enable quick, complete pipeline runs during development. Tools and pipelines for WGS can have a total CPU-cost of several days or weeks, which makes it cost-prohibitive to run structured testing of many tools using many samples. Towards the same goal one unit has constructed a “super-sick” sample by constructing a genome from a number of patient samples, in which a large number of variants can be tested in a single run.

During benchmarking and validation of new releases of a bioinformatics pipeline natural attention is given to the detection of clinically relevant variants. It is acknowledged that a pipeline release may fail to capture clinically important variants in a recall-test of previously detected variants, and the decision to accept this depends on the overall improvement in pipeline performance. Additionally, overall recall rates for variant detection may decline upon updating the truth set to more better and more comprehensive sets. Most bioinformatics pipelines run on a compute cluster will yield minor differences in repeated runs due to stochastic variability, and this should be accounted for when assessing pipeline performance, just like tests for robustness should be made.

##### Textbox 2: Nomenclature of quality assessment and optimisation in development of bioinformatics tools and pipelines.

**Benchmarking** systematically evaluates tools and methods against standardised test sets and criteria, focusing on performance metrics and suitability. **Verification** ensures that methods and tools meet design specifications. **Validation** confirms their reliability and accuracy in practical applications, addressing challenges such as genetic variation complexity, sequencing biases etc. **Testing** involves applying criteria to assess functionality, while **success criteria** define the acceptable standards for performance, accuracy, and reliability.

#### Release Management

Tools should be evaluated based on speed, cost, user acceptance, and performance metrics like F-scores, precision, and accuracy. Success criteria for pipeline changes need to be defined in advance to ensure changes achieve intended results. Version control and software release management is pivotal to ensure transparency and reproducibility of validation results. There are different structures of management of these processes in the different teams, but all teams report to work in close collaboration with clinical personnel to devise and improve the pipeline. Changes to the pipeline are thus requested, scoped and devised in a multidisciplinary dialogue with inclusion of all relevant competences and professions, and neither the users of the results nor the teams responsible for producing them are passive, or act merely on external requests.

#### Pipeline testing

After scoping a change request or addition to the pipeline and benchmarking a broad selection of tools and solutions, including one or more rounds of validation and test, a release candidate can be devised. Since bioinformatic pipelines consist of a number of successive processing steps with one step depending on the previous errors and imprecisions will propagate, but also many practical aspects including data annotation, formatting conventions, file naming or hierarchy may greatly influence the compatibility of tools in a pipeline. Therefore, testing of pipelines involves a combination of testing of individual components, testing integration, and end-to-end test of full pipeline. Below is an outline of typical steps that are needed, in parallel to biological and clinical understanding and evaluation.

**Unit Testing** typically involves use of a minimal datasets to allow rapid assessment of specific components within the pipeline. Synthetic data may be used to test targeted functionalities, ensuring that each component operates as expected. This approach is particularly useful when integrating new tools, as it allows for focused evaluation before full pipeline deployment, and a way to perform structured benchmarking of relevant tools. At this point in the development process there would often exist a “code freeze” or “release candidate” with a named git commit to be further evaluated. Any new code in the release candidate should be subject to manual code review.

**Integration Testing** involves running the entire pipeline with more extensive datasets to ensure comprehensive functionality and performance. It is recommended to use both minimal and larger datasets during this phase. For instance, a typical practice is to run patient samples through the pipeline to verify it meets predefined success criteria. Some groups also employ test-driven development methodologies, where tests are defined before making changes to the pipeline, ensuring that modifications achieve the desired outcomes.

**System testing and end-to-end testing** is performed at most sites (10/13) and includes verification and validation of the consequences on overall performance of the pipeline using representative datasets.

**IT performance testing** is evaluation of the system performance and responsiveness under different workloads and scenarios. It is often performed as stress-tests, e.g. by concurrent starting of many analyses pipelines. Additionally, security impact should be assessed and tested if specifically relevant to a release.

When to perform which test depends on the scope of the addition or change. As a general rule we suggest at major releases to perform end-to-end test, system test, performance and usability testing and at minor release to perform unit and integration testing, followed by regression and end-to-end testing. It will, however, depend on the change scenario and the internal definition of major or minor releases. The latter is often influenced by the assessment of the needed scale of testing.

#### Transition from Validation to Production

Transitioning from validation to clinical production requires careful management to ensure that the pipeline is ready for routine use. There was no consensus on whether to continue using minimal datasets or to employ larger, more representative datasets for this transition. One approach involves running actual patient samples and confirming that results align with expected results from previous pipeline versions or orthogonal testing. Manual interpretation is often still required in many cases, although efforts are being made to develop automated validation processes to reduce the workload on clinical staff. The workload associated with performing the manual validation is recognized as substantial - also to personnel outside of the bioinformatics team - and there is a clear need to automate as much of the process as possible to enhance efficiency. Nonetheless, manual review of output by interpretation specialists remains a critical component, particularly in cases where automated validation does not cover all aspects of pipeline performance. One unit reports the use of Continuous Integration testing (CI), where automated test scenarios are run upon each commit, merge and release.

##### Textbox 3: Key quality aspects in clinical bioinformatics production

**Use of established tools:** Utilise well-established and validated tools and algorithms.

**Accuracy:** Validate pipelines and implement quality control measures to detect and correct errors, and assess the performance.

**Reproducibility:** Ensure pipelines produce consistent results when run on the same data.

**Documentation and transparency:** Provide thorough documentation and transparency for understanding and troubleshooting.

**Specific considerations for different pipelines:** Address unique safety and quality requirements that are different e.g. for personalised treatment identification in cancer and rare disease diagnostics.

**Competence:** Employ experienced bioinformaticians to develop, review, upgrade, and maintain pipelines.

**Infrastructure:** Use suitable computational infrastructure to ensure scalability and reliability.

**External controls:** Participate in external quality assessment programs to benchmark pipeline performance.

**Figure 3.**
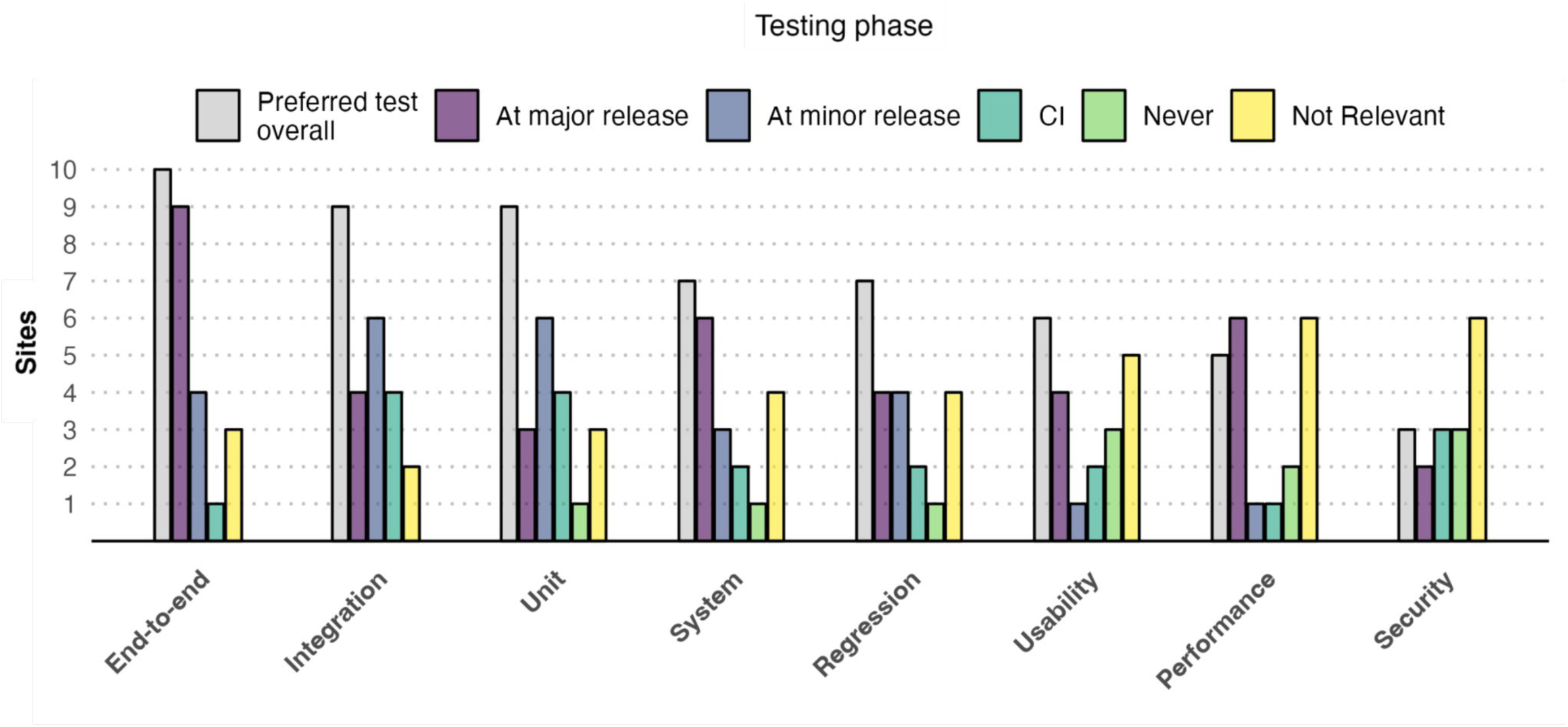
Reported use of test concepts to ensure quality of pipelines. The typical types of tests and test phases are shown on a polar barplot with the following notation for testing: Unit, testing individual components or functions in isolation; Integration, testing the interaction and integration between multiple components or modules; System, testing the entire system as a whole to ensure it meets the specified requirements; Regression, re-testing previously working functionality to ensure that changes have not introduced new issues; End-to-end, verifying the flow and functionality of the entire system from start to finish; Performance, evaluating the system’s performance and responsiveness under different workloads and scenarios; Security, identifying vulnerabilities and ensuring the system’s resistance against potential security threats; Usability, assessing the system’s user-friendliness and ease of use. The preferred test type overall is shown in grey. Counts on the barplot indicate the number of sites using each test type at each stage.

### IT management

#### High Performance Cluster

All teams agree that direct access to a High Performance Cluster (HPC) is recommended. Local HPCs can be preferable for security and availability, and ease of systems integration. Administration of such a cluster is a specialised task, that requires IT-competences that are not a priori part of a bioinformatician’s training. All teams have some tasks in terms of IT-management, but there are large differences on whether the main responsibility, and workload, is inside or outside of the team. Genomic information is per definition person-identifiable and sensitive information, and nearly all sites have a closed infrastructure not connected to the internet, exactly for security reasons.

#### Software containerisation and collaboration

One recommended route to robust production is to encapsulate software in containers (in production at respondents are Docker and Apptainer/Singularity), or alternatively, to predefine environments (in production at respondents are, Conda and EasyBuild) that the code should run in. This greatly reduces a number of risks that come from coexisting on shared IT infrastructures and it makes code sharing and co-development less dependent on where the code needs to run. Some have experienced sizable resource overhead from containerisation, which can be costly in a high-throughput production. The major obstacle is the magnitude of the data, which means that code has to be written to fit tight to the hardware for efficient and cost effective use of the cluster. This means that there are many parameters to balance at the expense of others, such as limitations on RAM, diskspeed, local node disk size, and read-write capacity (IO) to central storage, number of cores, and job queuing time - each of which could prove a bottleneck in one cluster, but not in another.

There are, however, very successful community development of bioinformatics pipelines that are containerized with a high level of platform adaptability in order to be less sensitive to the computer environment. Notable is the popular nf-core (https://nf-co.re/, (Ewels et al., 2020)) to which several of the units are contributing. Nf-core has an organised release structure, code review and continuous integration tests. An additional benefit from multicenter collaborations that is used actively and strategically, to avoid loss of competence when staff leaves, and it also contributes to increased safety and quality since multiple institutes, each with slightly different competencies and experiences, are involved.

#### IT security

Security is often not specific to a pipeline release, but is a continuous work assisted by dedicated IT-security teams in the organisation where there are also competences to perform Data Protection Impact Assessment (DPIA) when needed. Two-factor authentication (2FA) for server/GitHub access is now an expected minimum and enhances access security and prevents unauthorised access. Most sites use an air-gapped HPC (without direct access to the internet) and this is a general recommendation, as it removes a sizable risk component.

#### Pipeline Orchestration

Most teams use a workflow manager, most popular are Nextflow (4/8) or Snakemake (3/8), but also Cromwell and Stackstorm, and job schedulers like Crontab Dagu, are in production. A workflow manager allows for orchestrating parallelized workflows on a computer cluster where a bioinformatics pipeline can consist of multiple parts, each with different resource requirements and dependencies. Exceptions to this are two teams running commercial pipelines, one unit is using Nextflow to orchestrate jobs to Illumina DRAGEN hardware that can only process serially, and another team running the cloud based analysis service SOPHiA DDM by SOPHiA GENETICS. One team outsources the entirety of bioinformatics tasks to an external provider.

### Data and Metadata Management

Field standards for managing sequencing data and metadata in genomics informatics include ISO/TS 23357:2023 (ISO 23357, 2023), which outlines data fields from sequence reads to variant evaluation, and ISO/TS 20428:2024 (ISO 20428, 2024), which provides a template for clinical sequencing reports in electronic health record (EHR) systems. The HL7 FHIR (HL7 FHIR Foundation) standard facilitates effective data integration and transmission. Best practices involve using standardised formats, comprehensive metadata, and systems that enable smooth data exchange. Adhering to FAIR principles (Findable, Accessible, Interoperable, Reusable) and structured EHR templates is recommended for maintaining data quality and interoperability (Ryu et al., 2020).

Within NACG there is a high diversity in methods for approaching data and metadata management including flat files, sample sheets, gzip-compressed FASTQ files, various database technologies utilising some kind of common lab identifiers as well as clinical decision support software. Common practice is to use Laboratory Information Management Systems (LIMS) for these purposes. Integrations using the HL7 messaging standard are also worth mentioning (1/13).

In terms of propagation of the necessary metadata through the analytical pipelines, JSON and filenames are the most used methods (9/13 for both), closely followed by flat files (logs and other files; 7/13), database technology (e.g SQL, PostgreSQL; 6/13), yaml (5/13), environmental variables (4/13) and Nextflow channels (1/13). Most units use pseudonymization during data handling, to minimise availability of personal information.

There is no consensus on which files can be deleted from the production output, aside from SAM and BAM, that are now replaced by the compressed format CRAM. Some facilities save raw output from the machine (from Illumina these can have the file extension .bcl), whereas other’s discard these files after processing.

### Staff and competences for clinical production

The success of a clinical bioinformatics unit is highly dependent on the staff and their competencies. Key tasks include building data flows, setting up bioinformatic pipelines, developing front and backend applications, troubleshooting production runs, interpreting result quality, and ensuring regulatory compliance (NHS Health Education England, 2022).

The roles required vary significantly depending on the operation’s size and services offered. Smaller sites need staff with a broad skill set in order to fill the roles required, while larger facilities can afford more specialised positions. Essential competencies include software development, data management, quality assurance and domain knowledge in genetics. At larger sites with separate teams dedicated to system development, bioinformatic workflows or quality assurance, deep domain knowledge may be less critical for some positions, as teams can rely on each other’s expertise to fill the gap.

All sites agree that a highly automated workflow is preferable. Ideally, bioinformaticians should only need to troubleshoot results, with the system handling standard cases autonomously, from initiating workflows to delivering results. Automation enables staff to focus on more complex tasks and further refine the workflows and systems. Minimising repetitive tasks also helps keep staff engaged and motivated, which is crucial for retention.

Recruitment of new personnel is a challenge for most sites (12/13). The skills needed are often in high demand, making it difficult to find the right candidates. Traditionally, recruitment has been done through academic networks. However, as more specialised roles are required, it is necessary to look beyond these networks. Skilled front and backend developers, as well as system architects, are often not found in academic settings, making it necessary to advertise more broadly and possibly engage external recruiters. While most sites cannot match the salaries of the private sector, the NACG sites offer the chance to be part of a larger mission to the benefit of society, work with cutting edge technology, and participate in innovative research. Additionally, offering remote work options, flexible hours, and opportunities for internal training and career progression are strong motivators for many employees.

## List of figures

Figure SF1. Summary of provided analyses across units

Figure SF2. Data types used to validate and benchmark performance across tools

Figure SF3. Data and meta-data management

Figure SF4. Competences across NACG sites

## List of tables

Supplementary table ST1. Tools used in clinical production.

Supplementary table ST2. Datasets used to validate and benchmark (Summary list across all types of analysis)

