## Supplemental figures and tables for "Recommendations for Bioinformatics in Clinical Practice"

### Figures Supplementary

NACG paper

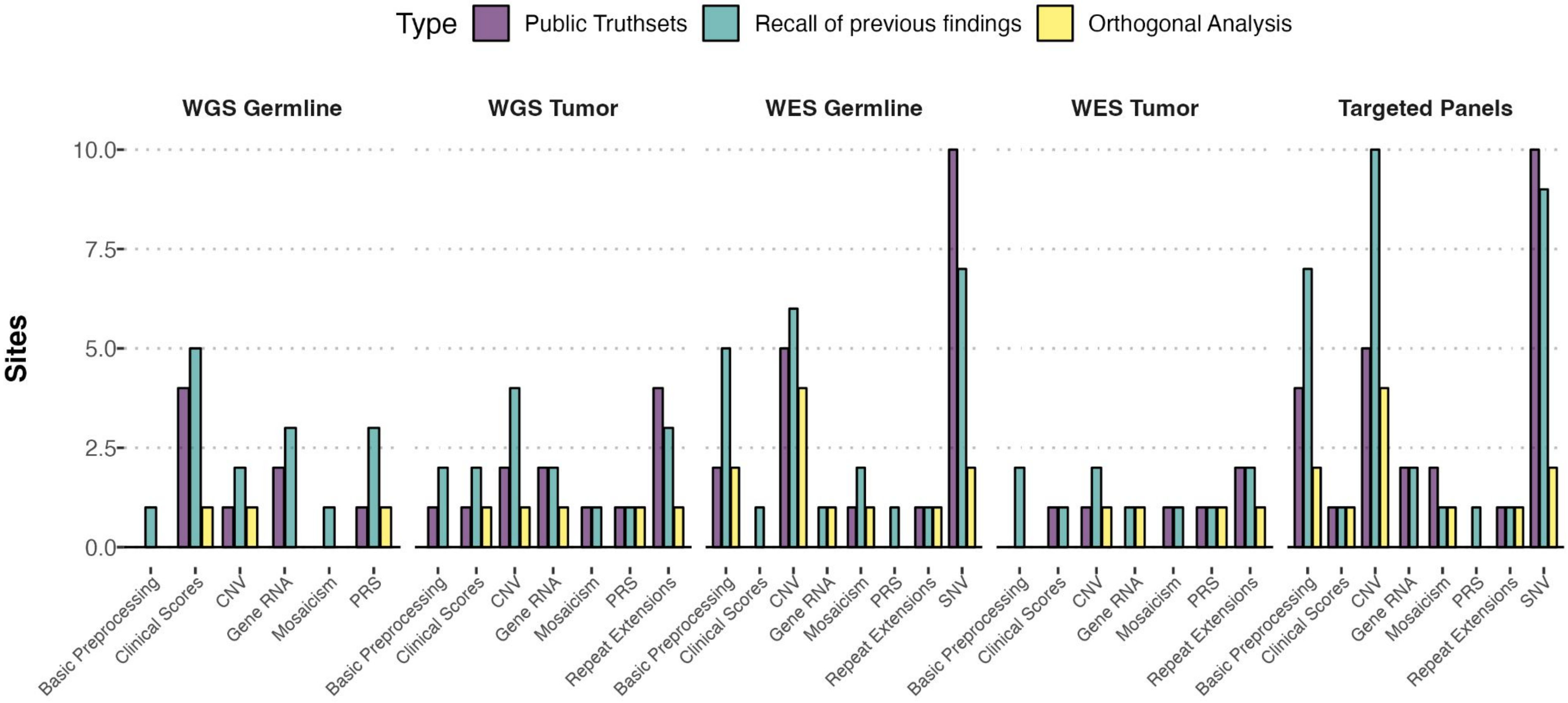

Type
Public Truthsets
Recall of previous findings
Orthogonal Analysis
WGS Germline
WGS Tumor
WES Germline
WES Tumor
Targeted Panels
Sites
10.0
7.5
5.0
2.5
0.0
Basic Preprocessing
Clinical Scores
CNV
Gene RNA
Mosaicism
PRS
Repeat Extensions
SNV

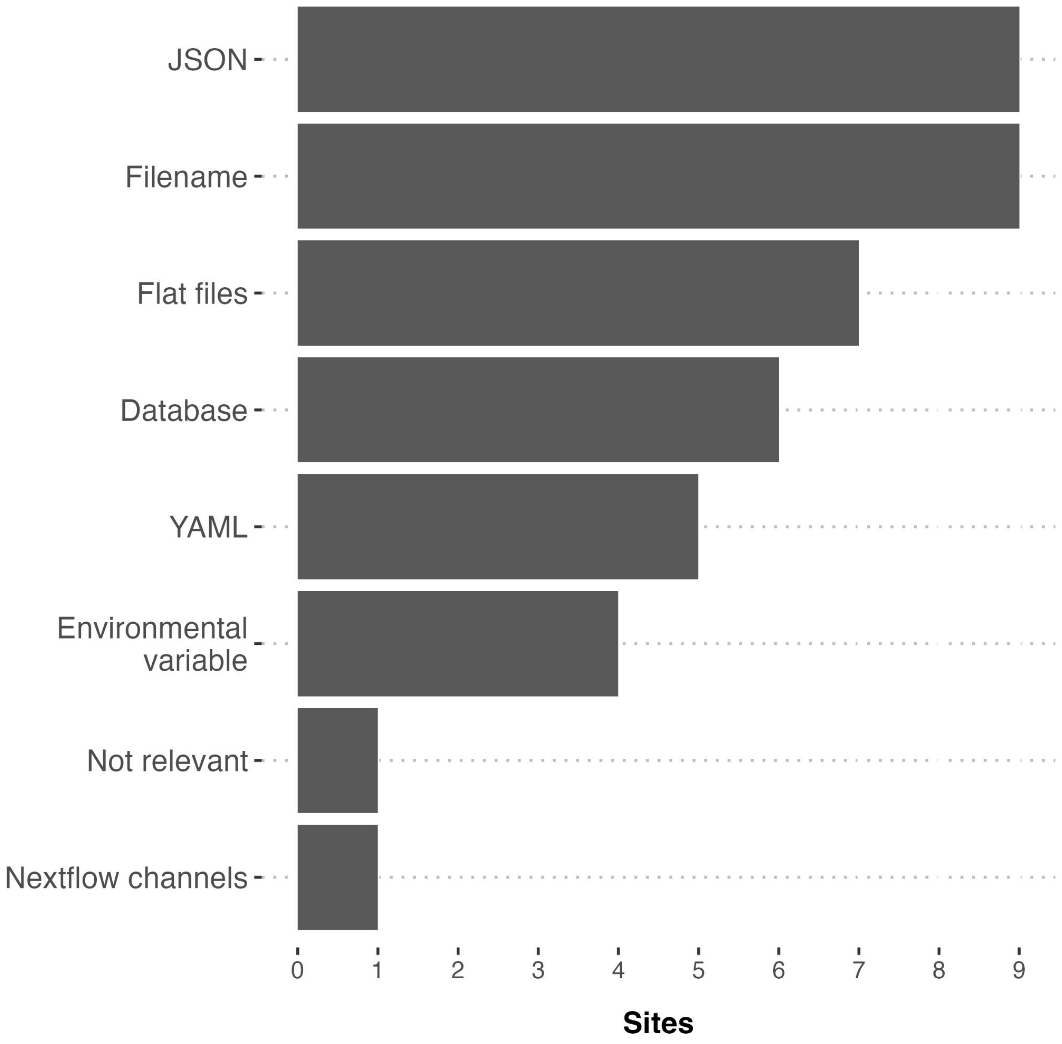


Figure SF2: Tools in clinical production to propagate metadata through the pipeline.

High diversity in methods for approaching data management

- csv files
- Sample sheets
- Flat files
- Gzip-Compressed Fastq-Files
- Laboratory Information Management Systems (LIMS)
- Multiple Databases (Common Lab IDs)
- HL7 Messaging Standard
- Ella and PostgreSQL-Based DBs

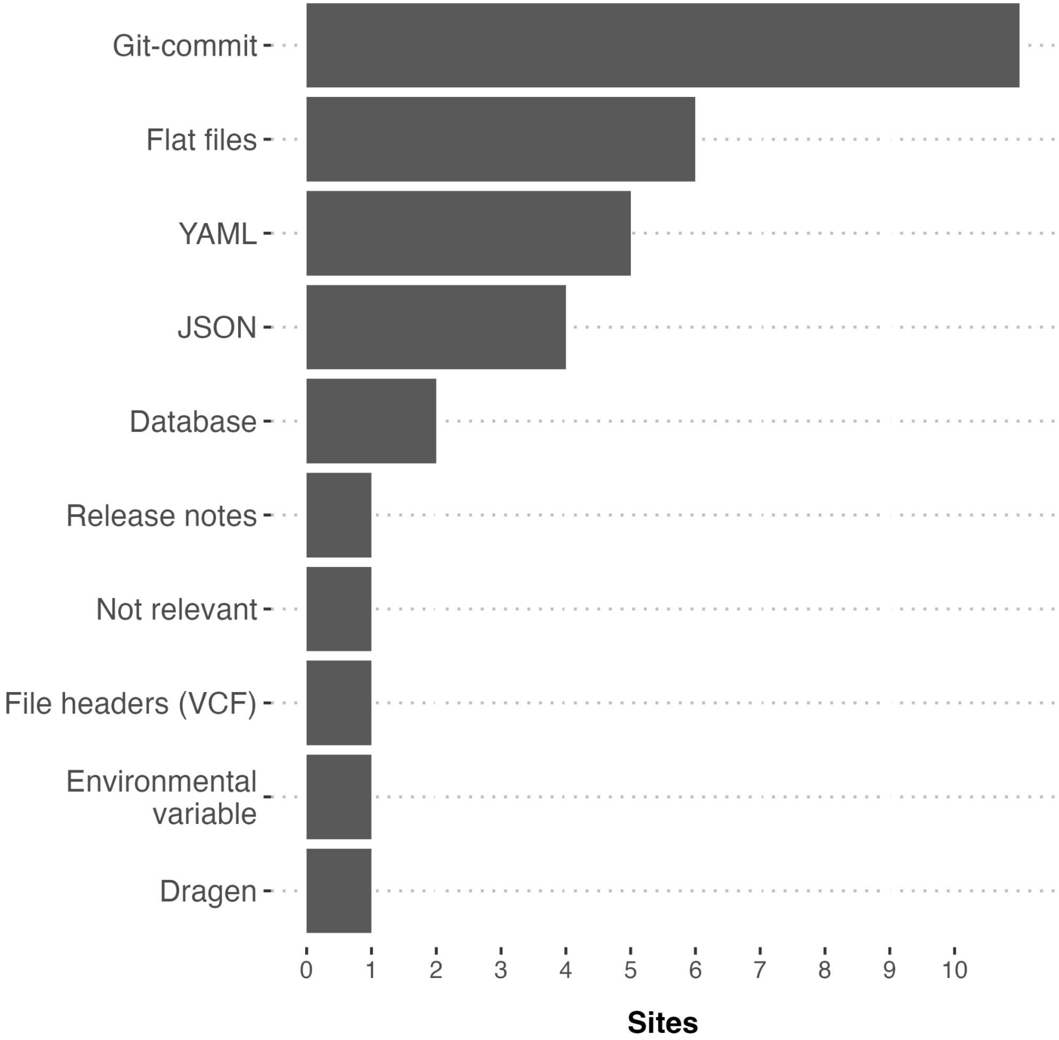


Figure SF3: Tools in clinical production for tracking version information for pipelines, databases and reference sets.

How to relate the output of each analysis to the version information

- VCF Files
- Log Files
  - Final reports
  - Snakemake Logs
- Configuration Files: YAML and JSON
- Git Commits
  - Flat files with Git-Commit
- Run Folder
- Sif Files
- Official Analysis Database

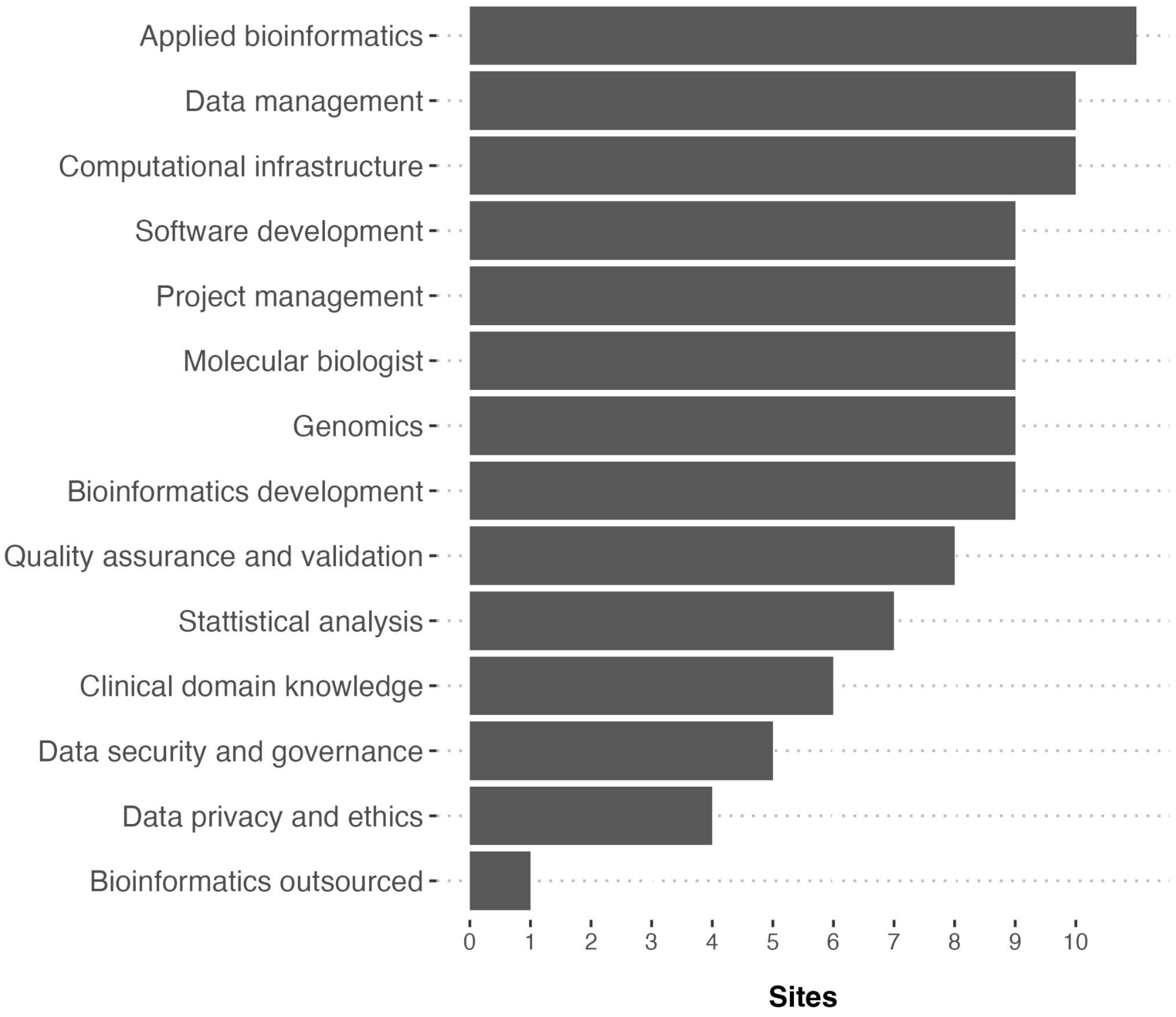


Figure SF4: Competencies needed for bioinformatic operations in a clinical context that are present in the team at each site. An individual can possess more than one competence.

#### Essential Competencies

- Knowledge of Bash/Linux/UNIX
- Proficiency in Python
- Familiarity with workflow languages
- Biostatistics skills
- Understanding of biology and genetics

#### Additional Competencies

- Version control
- Nextflow expertise (DSL1 and DSL2)
- Proficiency in Perl (for legacy code)
- Slurm management
- Data visualization using R-scripting
- Molecular biology knowledge, especially for cancer genomics
- Software engineering and stack development skills
- System administration and operation expertise
- Project management abilities
- Collaboration and communication skills
- Reporting capabilities

### Tables Supplementary

NACG paper

# ST1

Figure ST1: Bioinformatic tools commonly applied at various steps in clinical data analysis pipeline across sites. (p) denote the use of proprietary versions.

| | | |
|---|---|---|
| *Alignment* | DNA | bwa-mem, Sentieon (p), DRAGEN (p) |
| | RNA | STAR |
| *Small variant calling* | Germline variant | GATK, Sentieon (p), DRAGEN (p) |
| | Somatic mutation | Freebayes, Strelka2, Sentieon (p) |
| | Targeted panel | Freebayes, Pisces, VarDict, Sentieon (p) |
| *Large variant calling* | WGS | Manta, Tiddit, Lumpy, Delly, GATK, DRAGEN (p), cnvcaller |
| | WES | ExCopyDepth, ExomeDepth |
| | Targeted panel | CNVkit, CoNVaDING |
| | Other | OncoCNV (AmpliSeq), PureCN (TSO500) |
| *Custom* | Analysis | Fusioncatcher, Arriba, Archer Analysis (p) |
| *Variant* | Annotation | Nirvana, VEP, vcfanno, AnnotSV, Golden Helix varseq (p) |

# ST2

| | | |
|---|---|---|
| *Basic preprocessing* | WES | GIAB, Routine QC metrics, coverage, GA4GH |
| | WGS | GIAB, Routine QC metrics, Platinum genome, coverage |
| | Targeted panels | GIAB, Horizon discovery, Routine QC metrics, SeraSeq, coverage, Platinum genome |
| *Small Variant Calling* | Germline variant | GIAB (individual and trios, NA12878, hap.py,), Acrometrix, GA4GH, Platinum Genome |
| | Somatic mutation | GIAB, SEQC, Horizon discovery, Acrometrix, tool comparison |
| | Targeted panel | GIAB (hap.py, RTG tool), Horizon discovery, GA4GH, SeraCare (for oncogenic panels), Acrometrix, Platinum Genome |
| *Large variant Calling* | WGS | Previous cases and in-house comparisons, GIAB (HG002, TruVari) |
| | WES | Tool-comparison, GIAB (HG002), orthogonal analysis |
| | Targeted panel | GIAB, Horizon discovery, orthogonal analysis, In house comparisons, Acrometrix, SeraSeq |
| | Other | No benchmarking |
| *Gene Fusions* | WGS | Mini-gene, previous cases |
| | WES | Mini-gene |
| | Targeted panel | Horizon discovery, mini-gene, previous cases |
| *Mosaicism* | WGS | Previous cases (low VAF), in house controls, dilution experiment |
| | WES | Previous cases, Manual inspection of alignment and disease history. |
| | Targeted panels | Horizon Discovery (Mutect2), manual inspection. |
| *Repeat Extensions* | WES/WGS/Targeted Panels | Comparison to long-read sequencing and use positives to validate |

Table ST2: Recommended data to validate and benchmark performance. Note that quantification of the free answer text results generates implicit missing data and bias since not all participants was requested to take a stand on all data types.

# ST2

PRS: Only 1 reply "*This is not validated routinely. Run on a case by case basis and inspected manually.*" for targeted panels, WES/WGS germline

Clinical Scores:

- Targeted panels*:" Use of samples run by commercial applications such as myriad for HRD and comparing this to in-house developed software on the same samples. National collaborations sequencing the same samples and confirming results."*
- WES/WGS tumor*: compare different analysis to calibrate against known samples*
- WGS germline*: TMB; seracare samples, HDR; known braca1 cases, MSI previous case, Mutational signatures reported relative to cohort*
